# Phylogenetic relatedness of plant species co-occurring with an invasive alien plant species (*Anthemis cotula* L.) varies with elevation

**DOI:** 10.1101/2023.03.10.532156

**Authors:** Afshana, Jesús N. Pinto-Ledezma, Zafar A. Reshi

**Affiliations:** Department of Botany University of Kashmir, Srinagar-190006, Jammu and Kashmir, India; Department of Ecology, Evolution and Behavior, University of Minnesota, 1479 Gortner Ave, Saint Paul, MN, 55108, USA

**Keywords:** Darwin’s naturalization conundrum, Focal species, Phylogenetic metrics

## Abstract

Darwin’s naturalization conundrum, which posits that the alien species either succeed in the introduced region because being phylogenetically related to the native species hence being pre-adapted, or are phylogenetically dissimilar to native species and thus occupy unfilled niches, has received a lot of attention but the results have been contradictory. Instead of the usual phylogenetic comparison between native and introduced species to address this conundrum, we followed a novel approach of studying the phylogenetic relationship of a highly widespread and invasive species, Anthemis cotula L. (focal species) separately with the native species and all its co-occurring species (including native and non-native species) along an elevation gradient. The abundance of A. cotula declined continuously with an increase in elevation and species richness. The phylogenetic relationship between the focal species and all the co-occurring species using abundance-weighted mean pair-wise distance (MPDaw) showed an increase with an increase in elevation and species richness. A similar but slightly weaker relationship was noticed when the non-abundance weighted mean pair-wise distance (MPDpa) was used. Interestingly, the phylogenetic distance between the focal species and the native species based on MPDaw declined with elevation as well as species richness, but such a decline was seen when MPDpa was used. Our study also revealed that soil nutrients influence the abundance of A. cotula and the phylogenetic distance between the focal and other species, thereby indicating the role of micro-ecological factors and spatial heterogeneity in community assembly.

## Introduction

Invasive alien species are an outcome of a multi-stage process that begins with the intentional or unintentional transport and introduction of propagules of species by human beings into non-native regions well beyond their natural dispersal limits (Blackburn et al. 2011, Elton 2020). Humans have been transporting species around the globe for centuries and it has been accelerating in recent years due to an increase in trade and travel across countries and continents (Seebens et al. 2017). Today non-native species occur in Earth’s all continents and seas (Turbelin et al. 2017), in every biome, habitat, community and ecosystem (Elton 2020). While most of the alien species are benign, the highly invasive species impact community structure and functioning (Hejda et al. 2009; Linders et al. 2019), alter species interactions (David et al. 2017) or displace populations of native species (Pyšek et al. 2020). These impacts cause huge economic and ecological costs with enormous consequences for human well-being and livelihoods (Paini et al. 2016; Walsh et al. 2016; IPBES 2019). The damage costs from invasions stand at a staggering figure of US$1130.6 billion (Cuthbert et al. 2022) and the reported expenditures for the management of invasive species since 1960 have been estimated at about US$95.3 billion (in 2017). The impacts and costs due to these harmful invasive alien species are predicted to increase over the next decades worldwide (Essl et al. 2020; Seebens et al. 2021; Diagne et al. 2021).

What makes these alien species successful in the non-native ranges is one of the central issues in invasion biology, and many hypotheses have been put forth to explain the establishment and spread of non-native species. A recent review (Enders et al. 2020) reported at least 39 hypotheses that invoke one or the other explanation for the successful spread of alien species ranging from propagule pressure, enemy release, increased resource availability, and novel weapons, to empty niches hypothesis. This search for new and novel explanations is still continuing and new hypotheses are cited to explain the success of alien species, despite the bottlenecks and founder effects that can reduce genetic diversity initially. In this regard, Darwin also proposed two competing hypotheses (Darwin 1859), namely Darwin’s Naturalization hypothesis (DNH; Rejmánek 1996) and Pre-adaptation hypothesis (PAH; Ricciardi and Mottiar 2006). The Darwin’s Naturalization hypothesis suggests that alien species that are more distantly related to native species in the introduced range are more likely to succeed because they are able to exploit the underutilized niches, thereby minimizing competition and have fewer common natural enemies (Cadotte et al. 2018; Sheppard et al. 2018). Conversely, the pre-adaptation hypothesis, proposes that alien species that are closely related to native species in the introduced range are more likely to succeed because they share similar traits and adaptations that allow them to tolerate that environment and overcome environmental filtering (Mack 2003). The apparent contradition between the two hypotheses is known as Darwin’s Naturalization Conundrum (DNC) (Daehler 2001; Cadotte et al. 2018). While both hypotheses have some empirical support, the DNC remains a topic of debate among biologists. Indeed, the answer to this conundrum remains evasive because of the complex interactions and relative effects of ecological and evolutionary processes that drive the assembly of communities (Ng et al. 2018; Kusumuto et al. 2019; Park 2020; Pinto-Ledezma et al. 2020).

The attempts to resolve the Darwin’s Naturalization Conundrum have not been entirely successful, and how invasion success and phylogenetic distance between the native and introduced species are related has remained incosistent (Sol et al. 2022). This inconsistency has been attributed to scale-dependence of this relationship with Darwin’s naturalization hypothesis holding true at a small spatial scale while the pre-adaptation hypothesis being more applicable at a larger spatial scale (Park 2020, Qian and Sandel 2017). Put another way, the effect of abiotic and biotic factors on the community assembly and relationship between native and alien species varies at different spatial scales (Thuiller et al. 2010; Loiola et al. 2018; Pinto-Ledezma et al. 2020). Thus, alien species that are phylogenetically more closely related to the native species (phylogenetic clustering) are more likely to co-occur under conditions where abiotic factors have an overriding influence (Kraft et al. 2007; Leibold et al. 2010). Conversely, the alien species are phylogenetically distant to the native species (phylogenetic overdispersion) may co-occur in habitats where biotic competition structures the community assembly (Novotny et al. 2002; Webb et al. 2002; Godoy et al. 2014).

Recent evidence (Omer et al. 2022) showed that the relationship between invasion success and phylogenetic distance of introduced species to the native flora depended on the invasion stage. Introduction and invasiveness stages were positively related to phylogenetic distance, but converse was true for intervening naturalization process. This novel evidence highlights that the two apparently contradictory hypotheses of Darwin can act simultaneoulsy along the introduction-naturalization-invasion continuum (Omer et al. 2022). Other studies have brought out that changes in species interactions, such as facilitation, along a stress gradient may provide another explanation for DNC (Duarte et al. 2021; Wang et al. 2023) with Darwin’s naturalization hypothesis is supported under conditions of low stress and high interspecific competition; as a result, communities under these conditions are more resistant to invasion. Conversely, the preadaptation hypothesis is supported under high-stress conditions where the introduced species are more closely related to native communities and consequently more prone to invade local communities successfully.

Elevational gradients have been shown to have an important influence on the relative importance of biotic interactions and environmental filtering in determining community composition and invasion success (Duarte et al. 2021; Luo et al. 2023; McFadden et al. 2019). Studies have suggested that competition is more intense and frequent in low-elevation environments due to the relatively favorable environmental conditions that promote high species richness and overlap in resource use. In contrast, high-elevation environments characterized by harsher environmental conditions may limit the number of species that can survive and compete (Graham et al. 2012).

Based on this background information we studied the phylogenetic structure of plant assemblages at different elevations in Kashmir Himalaya that were invaded by *Anthemis cotula* L., a highly invasive ruderal species (Box 1). Specifically, we ask the following question: do the abundance, and phylogenetic relatedness between the focal invasive species, *A. cotula*, and the co-occurring native and alien plant species change with elevation? In addition, given that the number of competitors in local communities can influence the presence and abundance of invasive species—enemy release hypothesis (ERH; Keane and Crawley 2002)—we also asked: how does the number of species in local communities impact the abundance and phylogenetic structure of *A. cotula*?

We hypothesize that *A. cotula* would tend to be more associated with distantly-related species (phylogenetic overdispersion) at low elevations to escape stronger competition with closely-related species. Conversely, it should be in the company of closely-related species (phylogenetic clustering) at high elevations due to stronger environmental filtering. In addition, if the number of species that co-occur with *A. cotula* constrains its presence and abundance, we hypothesize a decrease in the focal species’ abundance with an increase in the species richness of local communities. We further hypothesize that the phylogenetic relatedness of *A. cotula* to co-occurring species would decrease with increasing species richness. Testing our hypotheses will shed light on the relative role of environmental and biotic factors in explaining the successful invasion of alien species.

### BOX 1. *Anthemis cotula* L. as focal species

*Anthemis cotula* L. is a monocarpic annual plant species that belongs to the family Asteraceae and is commonly known as ‘mayweed or stinking chamomile’, ‘dog fennel’ etc. The various synonyms of *A. cotula* are *Anthemis foetida* Lam., *Chamaemelum cotula* (L.) All., *Maruta cotula* (L.) DC., *Maruta foetida* Gray and *Matricaria cotula* (L.) Baill. (POWO, 2021). The species is native to Eurasia, especially in regions with a Mediterranean-type climate (Kay 1971; Erneberg, 1999) and Northern parts of Africa (POWO 2021). It is believed that this species was introduced outside its native range via trade as a contaminant of crop seeds/propagules (CABI 2019). *A. cotula*, owing to its ruderal life history strategy (r-selected), colonizes disturbed habitats and grows as a common weed in arable land, farmyards, roadsides, moist meadows, and overgrazed pastures. A syndrome of traits, such as synchronous germination of its achenes with favourable climatic conditions (Reshi et al. 2012), protracted recruitment and demographic trade-off with pre-winter and post-winter cohorts (Reshi et al. 2012), over-compensatory growth upon herbivory (Shah et al. 2012), allelopathic potential (Allaie et al. 2006; Rashid and Reshi 2012), widespread mycorrhizal association (Shah and Reshi 2007), high reproductive output (Kay 1971; Rashid et al. 2007) contribute to its invasiveness in Kashmir Himalaya. This rapidly spreading species is considered a threat to the native biodiversity (Adhikari et al. 2020) and this threat due to alien species is likely to exacerbate due to an increase in anthropogenic activities and global climate change (Richardson and Pyšek 2012; Downey and Richardson 2016).

## Methodology

### Study area

The study area for the present investigation was the Kashmir Himalaya which includes the main Kashmir Valley and also the side valleys of Tilel, Guraiz, Keran and Karnah (Dar and Khuroo 2013). The region is not only part of the biogeographic zone of the North-Western Himalaya in India (Rodgers and Panwar 1988) but also the Himalayan biodiversity hotspot (Dar and Khuroo 2020). It lies between geographical coordinates of 33° 20′ to 34° 54′ N latitude and 73° 55′ to 75° 35′ E longitude with an area of 15948 sq. km of which nearly 64% is mountainous (Khuroo et al. 2007). Topographically, the region mainly comprises a deep elliptical bowl-shaped valley which in the south and south-west is bound by the Pir Panjal range of Lesser Himalaya and in the north and north-east by the Zanskar range of the Greater Himalaya. This Valley of Kashmir is surrounded by high mountain ranges with the altitude of the main valley ranging from 1500 m to 1800 m (amsl), whereas the average height of its surrounding mountain ranges varies from 3000 to 4000 m, the highest peak being Kolahoi (5420 m). A characteristic and prominent geological feature of the region is the ‘Karawas’, which are plateau-like tablelands formed during the Pleistocene Ice age and are composed of clay, sand and silt of lacustrine origin (de Terra 1934). The valley is traversed by the river Jehlum and its tributaries which feed many world-famous freshwater lakes, such as the Wular, Dal and Anchar lakes. The climate of the region, marked by well-defined seasonality, resembles that of mountainous and continental areas of the temperate latitudes. The temperature ranges from an average daily maximum of 31°C and a minimum of 15 °C during summer to an average daily maximum of 4 °C and a minimum of – 4 °C during winter. It receives annual precipitation of about 1050 mm, mostly in the form of snow during the winter months. Owing to the vast variety of edapho-climatic and physiographic heterogeneity, the region harbours diverse habitats, including forests, grasslands, subalpine and alpine meadows, lakes, springs, swamps, marshes, rivers, cultivated fields, orchards, montane scree slopes and terraces, permanent glaciers etc., which support equally diverse floristic elements (Gupta 1982).

Like other parts of the world, Kashmir Himalaya is also beset with the scourge of alien plant invasions. A total of 571 vascular alien plant species, belonging to 352 genera and 104 families have been reported from the Kashmir Himalaya (Khuroo et al. 2007). The number of alien plant species was higher in families such as Poaceae (60 species), Asteraceae (54 species), and Brassicaceae (30 species). However, families such as Amaranthaceae (83%) and Chenopodiaceae (71%) showed a higher percentage of aliens relative to the total number of plant species belonging to these families in the region. Most of the alien plant species In Kashmir Himalaya (38%) were native to Europe followed by Asia (27%) and Africa (15%).

### Field sampling

Field surveys were conducted across the Kashmir Himalaya in 2018 and *A. cotula* was found growing over an altitudinal range of 1587 m (amsl) to 3700 m (amsl) (Fig. 1). Based on the population characteristics, particularly the size, 17 sampling sites were selected (each possessing >50% *A. cotula* cover) to cover more or less the entire spatial and altitudinal expanse of the species in the Kashmir Himalaya. The sampling sites supporting the populations of *A. cotula* were separated at least 15 km from each other. Though *A. cotula* was found to grow beyond 2700 m (amsl), such populations were not considered for phytosociological studies because these populations were comprised of a few scattered individuals. The geographic coordinates, elevation and soil characteristics of the sites harbouring the 17 populations are given in Table S1.

**Fig. 1.**
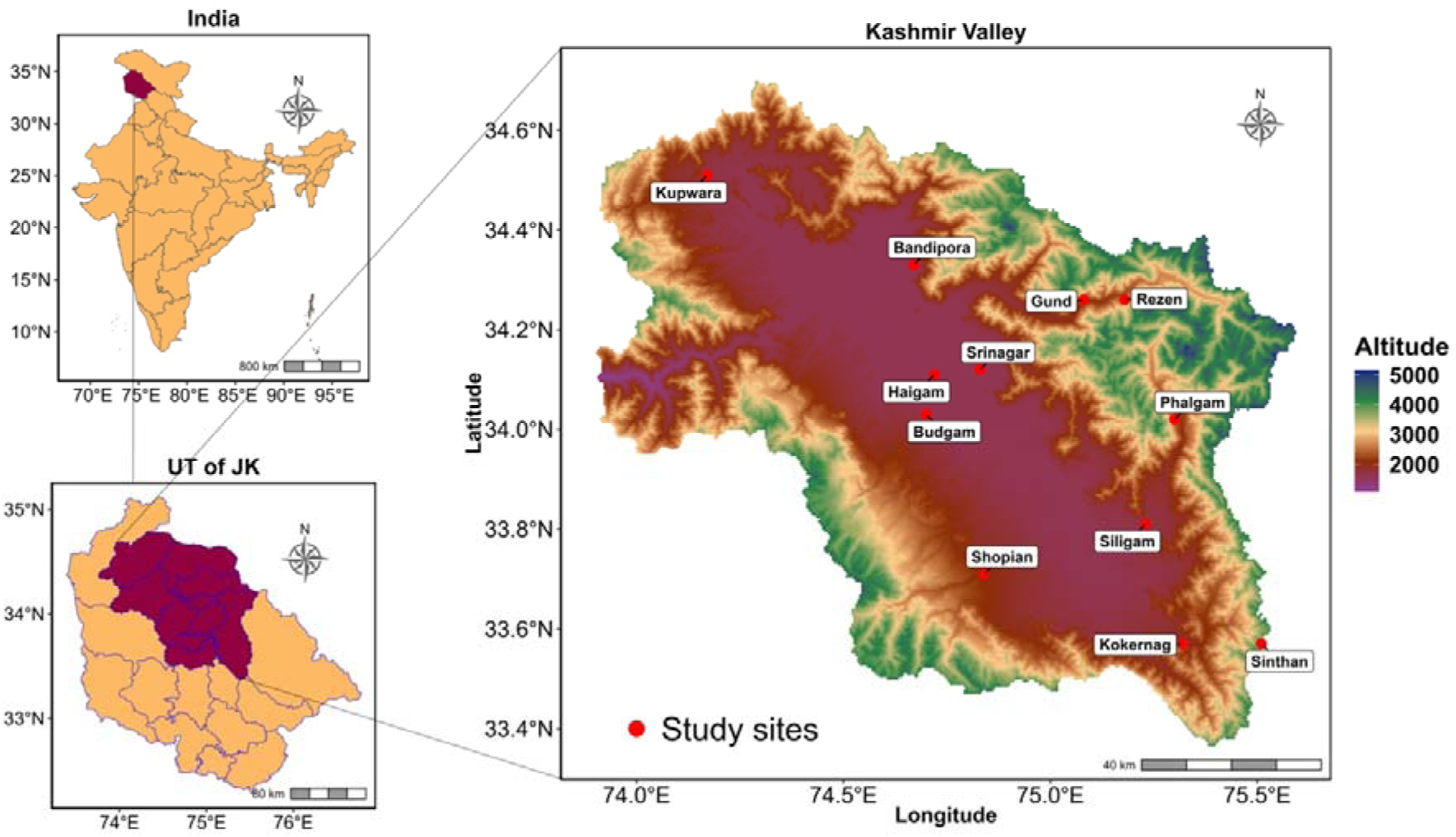
Location map of the study area and study sites.

**Table 1.** Taxonomic conspectus of taxa growing in association with *Anthemis cotula*

| Group | Species | Genera | Families |
| --- | --- | --- | --- |
| **Dicots** | 39 | 33 | 20 |
| **Monocots** | 6 | 6 | 2 |
| **Natives** | 30 | 27 | 19 |
| **Non-natives** | 15 | 13 | 98 |

A total of 10 quadrats (100×100 cm^2^) were laid randomly at each sampling site during the peak growing season. In each quadrat, all the plant species co-occurring with *A. cotula* were noted and their abundance was also recorded. The samples of plant species collected during the field survey were processed using routine herbarium techniques and identified using relevant literature, and herbarium records available in the Kashmir University Herbarium (KASH) and the help of experts was also sought wherever required. The synonyms and plant name authorships were ascertained from The Plant List (2013) using the function TPL() of the Taxonstand (Cayuela et al. 2012) package in R (R Core Team 2022). Moreover, various sources like, internet web pages, websites (www.efloras.org; https://powo.science.kew.org), Khuroo et al. (2007) were used to assign the nativity to each plant species. Based on life form, plant species were categorised into annuals, biennials, perennials and monocots and dicots as well.

### Phylogenetic tree

A phylogenetic tree (Fig. 2) was constructed using the R package V.PhyloMaker (Jin and Qian 2019). V.PhyloMaker uses the “GBOTB_extended.tree” as a backbone to insert missing species to their respective genera and families. This extended tree is an updated version of the Spermatophyta mega-phylogeny GBOTB (Smith and Brown 2018) and includes 74,533 species and all families of extant vascular plants. Branch lengths of the imputed species were set using the BLADJ approach as implemented in V.PhyloMaker (Jin and Qian 2019).

**Fig. 2.**
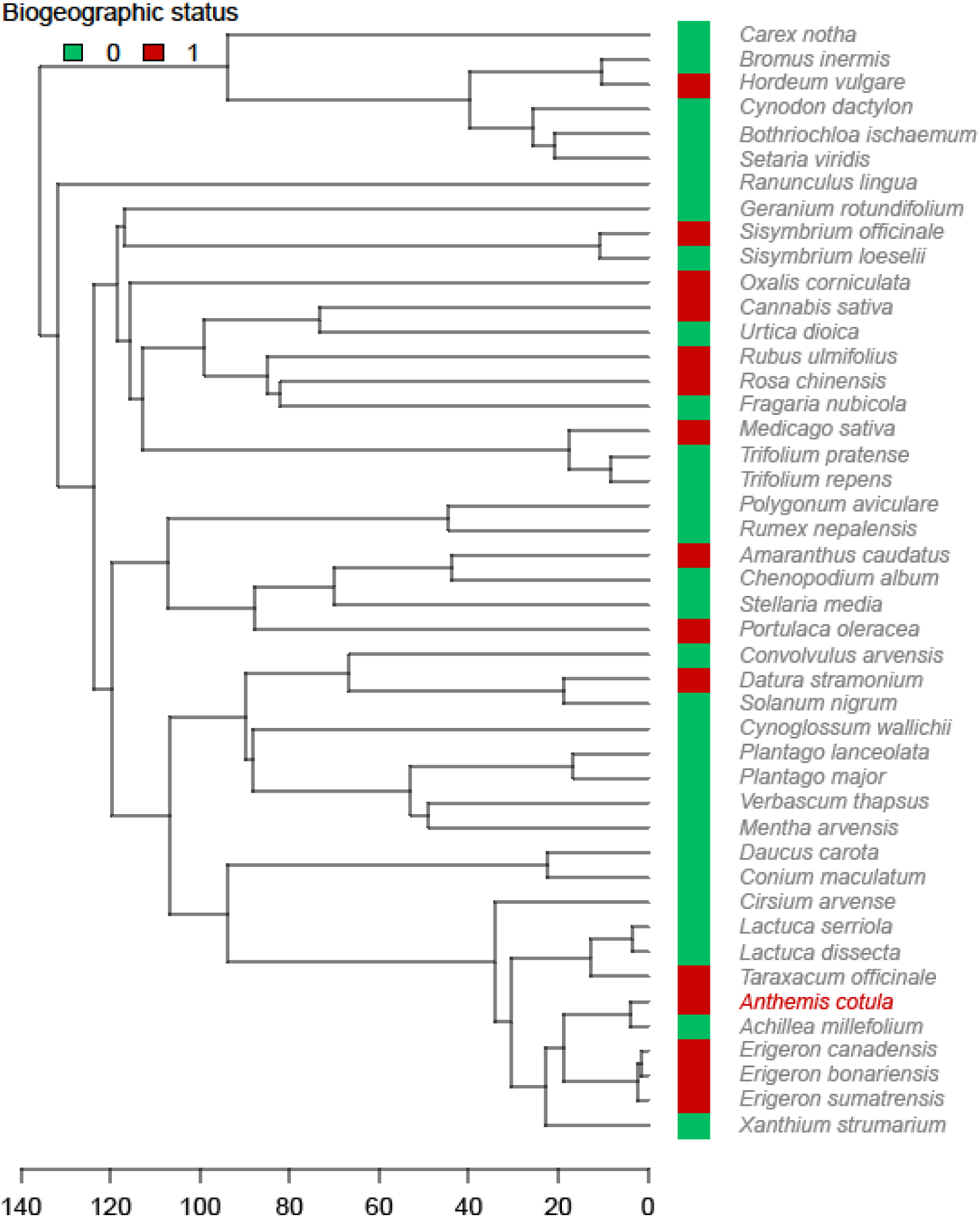
Phylogenetic tree used in this study. Green and red colors indicate the biogeographic status (native = 0 and introduced = 1). The focal-species (*Anthemis cotula*) is highlighted in red.

### Soil Analysis

A composite soil, representing a mixture of subsamples, was collected from each sampling site. Soil samples were collected by inserting a soil corer (2.25 cm diameter) vertically, and not tilted sideways, into the soil up to a depth of 15 cm. Five such soil samples from each site were thoroughly mixed and collected in plastic bags. In the laboratory, soil samples were gently crushed manually, air-dried and sieved through a 2-mm sieve to eliminate debris. The soil samples were then immediately processed for analysis.

The soil pH was measured for soil and distilled water suspension in the ratio of 5:1 using a portable pH meter (SYSTRONICS Model: MKVI). Organic carbon was determined using Walkley and Black’s (1934) rapid titration method. Olsen’s method (Olsen et al., 1954) was used for measuring phosphorus concentration. The flame photometry procedure outlined by Jackson (1973) was used to estimate total potassium (K) and sulphur was estimated by the calcium chloride method (Chesnin and Yien 1950). Calcium (Ca) was estimated in ammonium acetate extracts of soil by titration with EDTA (Cheng and Bray 1951). Soil characteristics are given in Table S1.

### Calculation of diversity metrics

To investigate the phylogenetic structure of *A. cotula*, we used the focal-species species approach (Pinto-Ledezma et al. 2020). The focal-species approach allows quantifying the phylogenetic structure of a set of species that co-occur with a focal species in a particular community or sampling site. Put simply, the focal species’ phylogenetic distance to all other community members is calculated and then averaged to obtain a single measure—e.g., MPD_focal_, mean phylogenetic-pairwise distance (MPD) of the focal species (see Pinto-Ledezma et al. 2020 for further details). We performed metric calculations that account for abundance (abundance-weighted MPD or MPD_aw_) and presence-absence (non-abundance-weighted MPD or MPD_pa_) of co-occurring species with *A. cotula* in each of the 17 sampling sites (Fig. 1).

Given that several non-native species were also found co-occurring with our focal species (Table 1), metric calculations were performed using all species (natives and non-natives) and only native species. Overall, we estimated 68 metric values (17 sampling sites x 2 metrics x 2 datasets) for *A. cotula*.

### Data analyses

Our analyses focused on assessing the probability of changes in abundance and phylogenetic relatedness of our focal species across elevation and local species richness. To do so, we modelled the association between the abundance and phylogenetic structure (MPD_aw_ and MPD_pa_) of our focal species against elevation and species richness using Bayesian linear models (BLMs). We assessed the effect of elevation or local species richness by testing the hypothesis that the high-density intervals (HDI) or 95% credible intervals (CI) of the BLM coefficients (slopes) do not overlap zero. To test the hypothesis for each model, we computed the evidence ratio (*ER*), which represents the posterior probability of a hypothesis against an alternative hypothesis—hypothesis = β ≠ 0, in other words, β is > 0 | < 0—where values of *ER* greater than one indicate evidence supporting the hypothesis.

In addition, given that additional environmental variables like soil properties may (or may not) influence the patterns of species co-occurrence, we additionally built models in which the abundance and the phylogenetic structure of *A. cotula* are influenced by the soil characteristics of the sampling sites. BLMs were implemented in the probabilistic programming language Stan (Carpenter et al. 2017) through the R package brms (Bürkner 2017). All analyses were run using 4 NUTS sampling chains for 5,000 generations and discarding 20% as burn-ins. Note that all BLMs were run twice, one using the all-species dataset (natives and non-natives) and the other using the only-natives dataset.

## Results

### General description of the data

In all 44 plant species were recorded in association with *A. cotula* belonging to 39 genera and 22 families (Table 1). Most of the species were dicots (39 species) and monocots were represented by only 6 species. Of the associated species, 30 were native to the region and 15 including *A. cotula* were non-native.

The number of species varied across sites with a minimum of 12 species recorded at the elevation of 1592 and 2666 m (amsl) and a maximum of 22 species recorded at an elevation of 2201 m (amsl) (Fig. S1). The native and non-native species belonging to dicots and monocots across sites is shown in Fig. S2. Number of species associated with *A. cotula* at the quadrat scale ranged from 6 to 9 across sites (Fig. S3).

A perusal of the data reveals that dicots were predominant across the elevations and the non-native species were entirely represented by dicots and no non-native monocot was recorded during the present study in any of the sites.

### Patterns of abundance and phylogenetic structure of A. cotula

The abundance of *A. cotula* varied considerably and showed a continued decline with increasing elevation and the number of species that co-occur in each sampling site (Fig. 3A-B). The pattern of continuous decline in abundance is consistent across datasets—i.e., using all species (natives and non-natives) dataset and the only native species dataset.

**Fig. 3.**
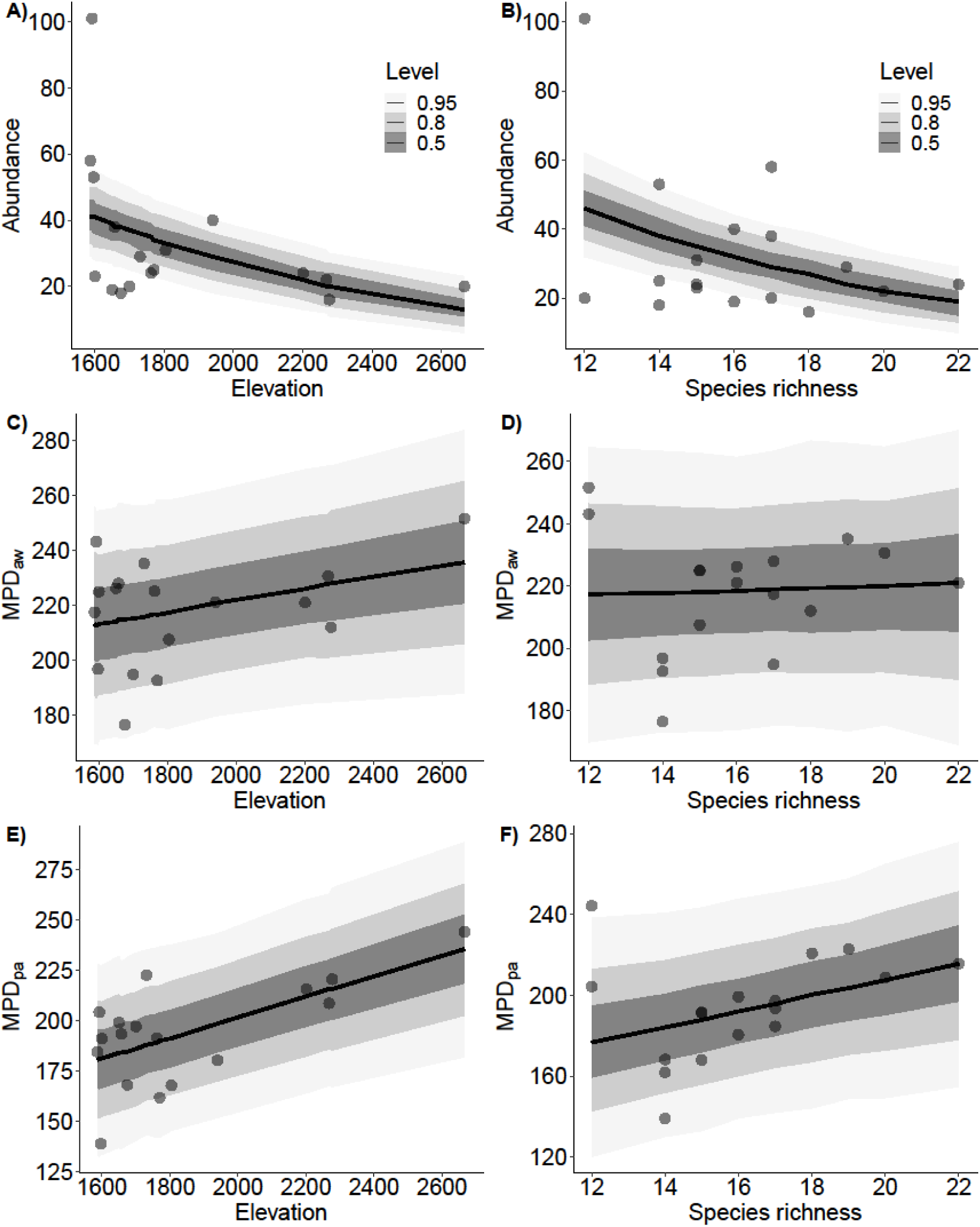
Marginal effects plots of changes in abundance and phylogenetic structure of *A. cotula* across elevation (lefthand panels) and local species richness (righthand panels) for the all-species dataset. Continuous black lines represent fitted slopes (with 50, 80, and 95% confidence intervals in gray).

Patterns of phylogenetic structure also varied, however, we observed shifts in the strength and direction of the associations between the degree of relatedness of *A. cotula* (measured as MPD_aw_ and MPD_pa_) and the covariables (elevation and richness). Specifically, we observed a shift from positive associations between MPD_aw_ and the covariables using the all-species dataset (Fig. 3C-D) to negative using the native-only dataset (Fig. S4C-D). For MPD_pa_, we found that the direction of the association remains but the strength decreases using the natives-only dataset (Fig. S4E-F).

### Effects of elevation and local species richness on patterns of abundance and co-occurrence of A. cotula

We found strong evidence that both elevation and local species richness influence negatively the abundance of *A. cotula* and that the effect is consistent across datasets (Fig. 4A-B). Indeed, our Bayesian models reveal strong evidence (*ER* = Inf; *ER*_natives-only_ = Inf) for the effect of both covariables on *A. cotula* abundance patterns (Table 2). For the phylogenetic structure under the all-species dataset we found strong evidence for the effect of elevation on MPD_pa_ (*ER* = 375.471) and small evidence for the effect local species richness on MPD_pa_ (*ER* = 15.285). For MPD_aw_ we found small evidence for the effect of elevation (*ER* = 11.346) and weak evidence for the effect of local species richness (*ER* = 1.346) (Fig. 4D, 4F). Under the natives-only dataset, we found small evidence for the effect of elevation on MPD_pa_ (*ER* = 8.44) and MPD_aw_ (*ER* = 2.507). Local species richness exerts a strong effect on the patterns of MPD_aw_ (*ER* = 425.667) but weakly on the patterns of MPD_pa_ (*ER* = 2.507) (Fig. 4C, 4E, Table 2).

**Fig. 4.**
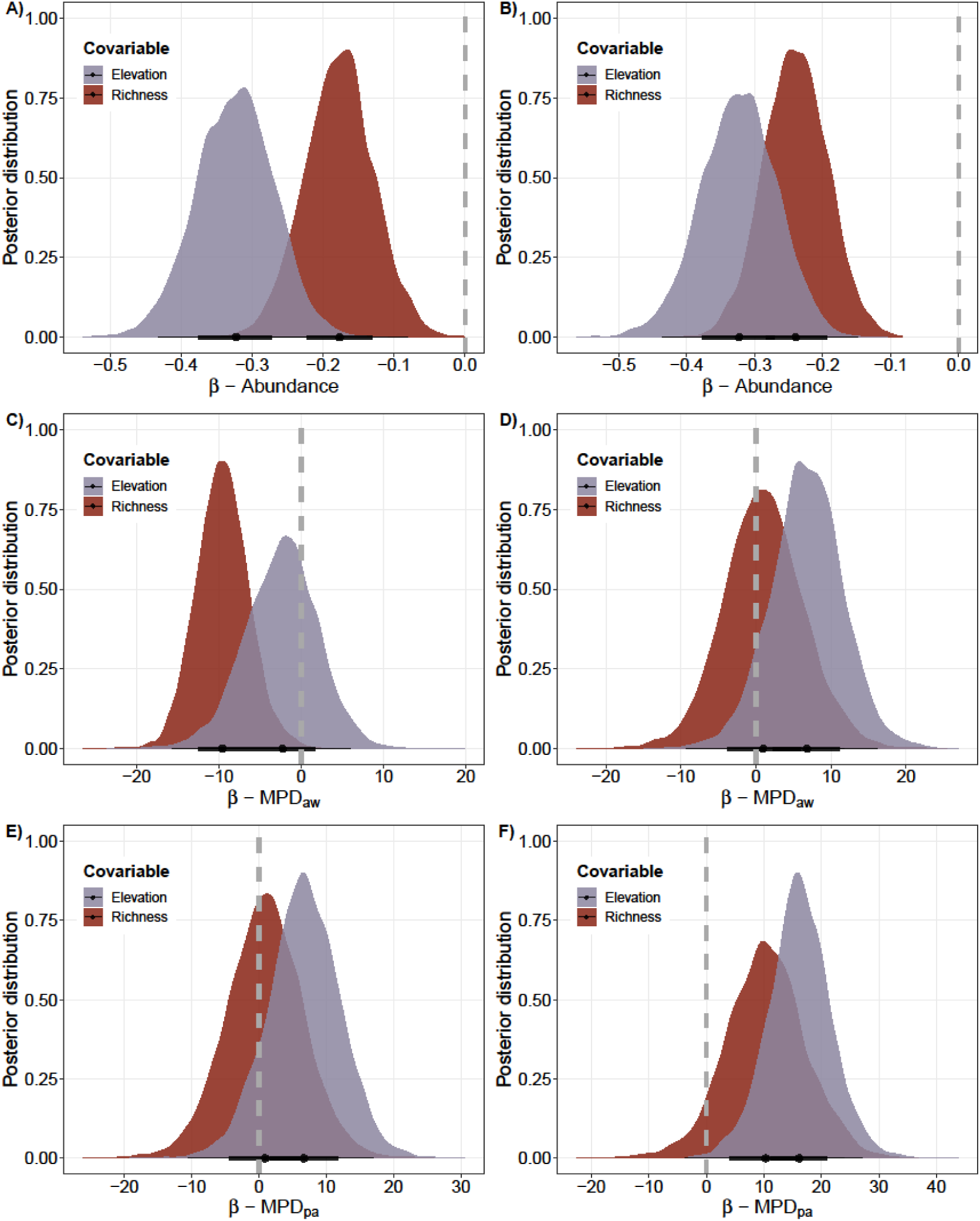
Posterior distribution of the marginal effects (slope coefficients [β]) of elevation and local species richness on *A. cotula* abundance and phylogenetic structure. Each point (shown with black bars) represents the median β with its associated 95% high-density credible interval (HDI). Lefthand panels (A, C, E) correspond to the natives-only dataset and the righthand panel (B, D, F) to the all-species dataset.

**Table 2.** Comparative hypothesis testing for the biotic (species richness) and abiotic (elevation) covariables influencing the abundance and the phylogenetic structure of our focal-species. Bold-faced rows indicate that the influence of the covariable is different from zero, i.e., strong evidence. See methods section for details of the Evidence Ratio (*ER*) calculation and interpretation.

| Dataset | Variable | Covariable | Estimate | CI Lower<br>-2.50% | CI Upper<br>97.50% | Evidence<br>Ratio | Posterior<br>Distribution | Star |
| --- | --- | --- | --- | --- | --- | --- | --- | --- |
| All species | Abundance | Elevation | **-0.323** | **-0.416** | **-0.234** | *Inf* | **1** | * |
| All species | Abundance | Richness | **-0.24** | **-0.315** | **-0.162** | *Inf* | **1** | * |
| All species | MPD<sub>aw</sub> | Elevation | 6.787 | -1.196 | 14.543 | 11.774 | 0.922 |  |
| All species | MPD<sub>aw</sub> | Richness | 0.955 | -7.513 | 9.962 | 1.346 | 0.574 |  |
| All species | MPD<sub>pa</sub> | Elevation | **16.108** | **7.21** | **25.097** | **375.471** | **0.997** | * |
| All species | MPD<sub>pa</sub> | Richness | 10.296 | -0.826 | 21.877 | 15.285 | 0.939 |  |
| Native only | Abundance | Elevation | **-0.323** | **-0.415** | **-0.237** | *Inf* | **1** | * |
| Native only | Abundance | Richness | **-0.177** | **-0.258** | **-0.098** | *Inf* | **1** | * |
| Native only | MPD<sub>aw</sub> | Elevation | -2.271 | -9.79 | 4.558 | 2.507 | 0.715 |  |
| Native only | MPD<sub>aw</sub> | Richness | **-9.598** | **-14.659** | **4.442** | **425.667** | **0.998** | * |
| Native only | MPD<sub>pa</sub> | Elevation | 6.68 | -2.27 | 15.323 | 8.44 | 0.894 |  |
| Native only | MPD<sub>pa</sub> | Richness | 0.909 | -8.525 | 10.137 | 1.31 | 0.567 |  |

We were also interested in exploring the effect of soil covariables on the patterns of abundance and phylogenetic structure (Fig. 5). We found strong evidence that SOC, S, K, and N influence the abundance of *A. cotula* (*ER*[64.844, Inf], Table S2). For the phylogenetic structure, it seems that soil covariables do not strongly influence the patterns of MPD_aw_ and MPD_pa_ for both datasets (Fig. 5B-E), however, we found strong evidence for the effect of Ca on MPD_aw_ (*ER*_all-species_ = 40.237, *ER*_natives-only_ = 40.885) (Fig. 5B-C). We also found strong evidence for the effect of SOC on MPD_pa_ for both datasets (*ER*_all-species_ = 64.844, *ER*_natives-only_ = 27.470) (Fig. 5D-E).

**Fig. 5.**
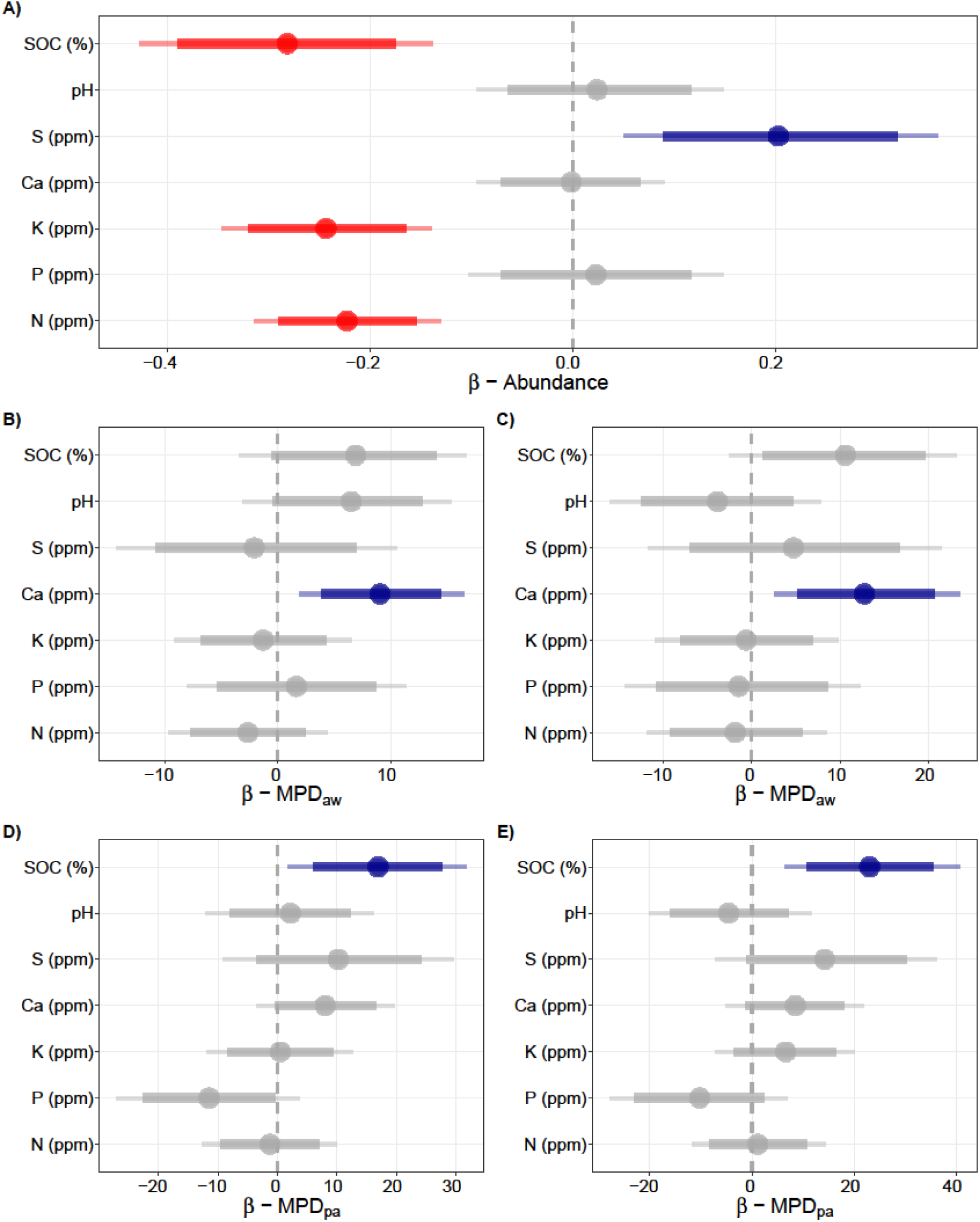
Marginal effects (slope coefficients [β]) of soil covariables on A. cotula abundance and phylogenetic structure. Each point represents the median β with its associated 89% and 95% high-density credible interval (HDI). Blue and red colors indicate positive and negative effects of soil covariables, respectively. Panels B and D correspond to the natives-only dataset and the panel C and E to the all-species dataset. Gray colors indicate no effect.

## Discussion

The co-occurrence patterns of the invasive *A. cotula* in invaded communities in the Kashmir Himalaya revealed that abiotic (elevation) and biotic (species richness) play contrasting roles in explaining its successful invasion. Our results revealed that the number of species co-occurring with *A. cotula* varied across spatial scales, i.e., at landscape, site and quadrat scales. We also found a consistent decline in the abundance of our focal-species with increasing species richness in local communities (Fig. 3B, S4B), thus supporting the enemy release hypothesis (ERH). Moreover, the patterns of phylogenetic structure change show a tendency to overdispersion (i.e., support for Darwin’s naturalization hypothesis (DNH)) with increasing elevation (Fig. 3C-E); however, this pattern is challenged using the natives-only dataset (Fig. S4C). Specifically, the patterns of phylogenetic structure strongly depend on the nature (i.e., abundance-weighted, presence-only) of the data used for metric calculations. Our findings provide novel insights for understanding the successful invasion of introduced species across abiotic and biotic gradients in the Himalaya, a region historically impacted by human activities.

Evaluating the number of co-occurring species with our focal-species (*Anthemis cotula* L.), we found that at the landscape scale, 44 species were co-occurring with *A. cotula,* while on average, only 15 and 6 species co-occurred at the site and quadrat scales, respectively (Figs S1-S3). It brings out that number of co-occurring species at the landscape scale is higher compared to the number of species at smaller spatial scales. It means that different plant species get associated with *A. cotula* across sites and hence a higher number of co-occurring species at the landscape scale. Variations in species richness at different spatial scales have been well-reported (Chase et al. 2019). While at a landscape scale, evolutionary processes, and topographical factors (e.g., elevation) generally are related to variation in species richness (McFadden et al. 2019; Pinto-Ledezma et al. 2018), at a local scale, processes, such as species interactions, disturbance, and grazing (Bhattarai 2017; Pinto-Ledezma et al. 2020), are known to influence species richness (Vetaas 1997). Thus, anthropogenic disturbances characteristic of ruderal habitats used by *A. cotula* may be creating spatial heterogeneity that structures the species association patterns (Gao et al., 2021). We also found a consistent decline in the abundance of *A. cotula* with increasing elevation (Fig. S5A). The decrease in abundance with elevation could be due to a reduced number of propagules of *A. cotula* and harsh climatic conditions in the higher elevations which may be preventing it from attaining higher abundance.

Our findings also revealed an increase in MPD_aw_ and MPD_pw_ with the elevation and richness of all species (native and non-native) (Fig. 3). In other words, increasing phylogenetic distances between *A. cotula* and all species pointed towards phylogenetic overdispersion with increasing elevation (Fig. 3C-E). This result contrasts our hypothesis and many previous studies that have reported the co-occurrence of phylogenetically closely related species (phylogenetic clustering) at higher elevations due to harsh and stressful conditions (e.g., González-Caro et al. 2014; Machac et al. 2011; Qian et al. 2013). Our finding of phylogenetic overdispersion with increasing elevation, however, draws support from similar results in the Rocky Mountains (Bryant et al. 2008), Andes (Qian 2018), and Malesia of tropical Asia (Culmsee and Leuschner 2013).

Being a species that thrives in disturbed habitats, we expected lower phylogenetic distances between *A. cotula* and other co-occurring species because of lower competition which favours phylogenetically distant species due to competitive exclusion (Slingsby and Verboom 2006; Webb et al. 2006) but the results are contrary to our expectations. A potential explanation for this pattern is that disturbance creates conditions suitable for *A. cotula* and other distantly-related co-occurring species. Also, it must be noted that *A. cotula* tracks roads and trails, which also serve as conduits for the spread of alien species, thereby increasing the phylogenetic distance between the species assemblages outweighing the effect of reduced competition. Conversely, MPD_aw_ between *A. cotula* and native species dataset declined with elevation and species richness depicting phylogenetic clustering at higher elevations and phylogenetic over-dispersion at lower elevations (Fig. S4C,D). This observation is in agreement with many previously reported similar findings (Webb et al. 2002; Li et al. 2014; Manish and Pandit 2018). Thus, it is quite apparent that the phylogenetic relationship of *A. cotula* varies with native and all species (including introduced and natives). This adds another dimension to the already complex phylogenetic relationship between introduced and native species. While comparing the phylogenetic distance between *A. cotula* and native species, our hypothesis of phylogenetic overdispersion at low elevations and phylogenetic clustering at high elevations is supported but the same is rejected when the comparison is between *A. cotula* and native and non-native species taken together.

The preceding discussion reveals that our hypothesis of phylogenetic overdispersion at low elevations—to escape stronger competition with closely related species—and phylogenetic clustering at high elevations—due to stronger environmental filtering—is supported when we studied the phylogenetic relationship between *A. cotula* and its co-occurring native species (Fig. S4C). Simply put, *A. cotula* tended to be associated with distantly-related species to escape stronger competition with closely-related native species. This trend is reversed when the phylogenetic relationship between *A. cotula* and its co-occurring native, as well as non-native species, was studied (Fig. 3C, 3E). It appears that microecological factors and not macroecological conditions (general climate) shape the species association patterns when native and non-native species are taken together, but in respect of native species, long-term evolutionary processes may determine the species’ co-occurrence with *A. cotula*. Indeed, it is well known that the microclimate of a plant species is highly heterogeneous across space and time and differs strongly from the surrounding macroclimate (Kearney and Porter 2009; Sears et al. 2011) and that anthropogenic disturbances may be contributing to this heterogeneity and hence creating empty niches for non-native species to occupy.

The effect of elevation and species richness was apparent when abundance-weighted MPD was used compared to presence-absence-weighted MPD (Fig. 4). Abundance-based measures detect more nuanced variation in the abundance of species across sites and it is quite relevant in respect of invasive species where abundance is of overriding importance and not the species identity alone (Cadotte et al. 2010; Barwell et al. 2015). In fact, species abundance is critical for ecological processes and dynamics (Hillebrand et al. 2008), and its inclusion in metric calculations allows the detection of changes in the species composition and phylogenetic structure of ecological communities (Miller et al. 2017; Pinto-Ledezma et al. 2020). Evidence of this observation was presented in simulation studies (e.g., Miller et al. 2017) and natural settings. For example, changes from phylogenetic clustering to random structure were found in the western Amazon basin (Eiserhardt et al. 2013), and changes from phylogenetic clustering to overdispersion in eastern Australia (Sommer et al. 2017).

At smaller scales, abiotic factors, such as soil characteristics are considered local drivers of the fine-scale richness patterns of non-native species in mountain ecosystems (Buri et al. 2017; Gantchoff et al. 2018; Lembrechts et al. 2019). Specifically, we found a negative effect of soil organic carbon, K and N contents, and a positive effect of soil sulphur content on the abundance of *A. cotula* (Fig. 5). Among all the soil variables, soil calcium positively influenced MPD_aw_ irrespective of whether native only or all species were considered. Likewise, soil organic carbon had a significant positive effect on MPD_pa_ but soil calcium and phosphorus had a significant negative effect on MPD_pa_ (Fig. 5). The relationship between species richness, species abundance and phylogenetic diversity, and soil nutrients along elevation gradients is difficult to discern, but increase in species and phylogenetic diversity with a decline in soil nutrients has been reported (Sander and Wardell-Johnson, 2011). Soils at higher elevations are usually nutrient-poorer, but the abundance of species is high with larger phylogenetic distances between species (Sander and Wardell-Johnson, 2011). Given complex interactions between biotic and abiotic factors that determine the availability of soil nutrients, the actual mechanisms as to how soil nutrients influence the phylogenetic structure at the ecological scale are difficult to distinguish. Further research is needed to fully understand the extent to which soil nutrients influence the successful invasion of introduced species. These efforts may contribute to addressing critical hurdles posed by species invasion and developing effective management strategies in the Anthropocene.

## Acknowledgements

We thank Head, Department of Botany, University of Kashmir for providing laboratory facilities. Award of fellowship by the UGC-CSIR, India to the corresponding author (Afshana) and the support under the CPEPA by the UGC, New Delhi to the University of Kashmir is also gratefully acknowledged. JPL was supported by the US National Science Foundation (DEB-2017843 to JPL).

## Data availability

The data could be made available by the corresponding author on reasonable request.

## Author’s contribution

Afshana and ZAR did sampling and generated the data, JPL and ZAR analyzed the data. ZAR, JPL and Afshana wrote and revised the manuscript. All authors have read and agreed with the final version of the manuscript.

## References

Adhikari S, Burke IC, Eigenbrode SD (2020) Mayweed chamomile (*Anthemis cotula* L.) biology and management—a review of an emerging global invader. Weed Res 60(5):313–322.

Allaie RR, Reshi Z, Rashid I, Wafai BA (2006) Effect of aqueous leaf leachate of *Anthemis cotula*–an alien invasive species on germination behaviour of some field crops. J Agron Crop Sci 192(3):186–191.

Barwell LJ, Isaac NJ, Kunin WE (2015) Measuring β[diversity with species abundance data. J Anim Ecol 84(4):1112–1122.

Bhattarai KR (2017) Variation of plant species richness at different spatial scales.BOTOR: J Plant Sci 11:49–62.

Blackburn TM, Pyšek P, Bacher S, Carlton JT, Duncan RP, Jarošík V et al (2011) A proposed unified framework for biological invasions. Trends Ecol Evol 26(7): 333–339.

Bryant JA, Lamanna C, Morlon H, Kerkhoff AJ, Enquist BJ, Green JL (2008) Microbes on mountainsides: contrasting elevational patterns of bacterial and plant diversity. Proc Natl Acad Sci 105(supplement_1):11505-11511.

Buri A, Cianfrani C, Pinto-Figueroa E, Yashiro E, Spangenberg JE, Adatte T et al (2017) Soil factors improve predictions of plant species distribution in a mountain environment. Prog Phys Geogr 41(6):703–722.

Bürkner PC (2017) brms: An R package for Bayesian multilevel models using Stan. J Stat Softw 80:1–28.

Cadotte MW, Campbell SE, Li SP, Sodhi DS, Mandrak NE (2018) Preadaptation and naturalization of nonnative species: Darwin’s two fundamental insights into species invasion. Ann Rev Plant Biol 661–684.

Cadotte MW, Jonathan Davies T, Regetz, J, Kembel SW, Cleland E, Oakley TH (2010) Phylogenetic diversity metrics for ecological communities: integrating species richness, abundance and evolutionary history. Ecol Lett 13(1):96–105.

Carpenter B, Gelman A, Hoffman MD, Lee D, Goodrich B, Betancourt M et al (2017) Stan: A probabilistic programming language. J Stat Softw 76(1).

Cavender-Bares J, Keen A, Miles B (2006) Phylogenetic structure of Floridian plant communities depends on taxonomic and spatial scale. Ecol 87(sp7):S109-S122.

Cayuela L, Granzow de la Cerda Í, Albuquerque FS, Golicher DJ (2012) Taxonstand: An R package for species names standardisation in vegetation databases. Methods Ecol Evol 3(6):1078–1083.

Chase JM, McGill BJ, Thompson PL, Antão LH, Bates AE, Blowes SA et al (2019) Species richness change across spatial scales. Oikos 128(8):1079–1091.

Cheng KL, Bray RH (1951) Determination of calcium and magnesium in soil and plant material. Soil Sci 72:449–458.

Chesnin L, Yien CH (1950) Turdimetric estimation of sulphates. In Soil Science Society of America (Vol. 15, pp. 149–151).

Culmsee H, Leuschner C (2013) Consistent patterns of elevational change in tree taxonomic and phylogenetic diversity across Malesian mountain forests. J Biogeogr 40(10): 1997–2010.

Cuthbert RN, Diagne C, Hudgins EJ, Turbelin A, Ahmed DA, Albert C et al (2022) Biological invasion costs reveal insufficient proactive management worldwide. Sci Total Environ 819:153404

Daehler CC (2001) Darwin’s naturalization hypothesis revisited. Am Nat 158(3):324–330.

Dar GH, Khuroo AA (Eds.) (2020) Biodiversity of the Himalaya: Jammu and Kashmir State (Vol. 18). Singapore. Springer.

Dar GH, Khuroo AA (2013) Floristic diversity in Kashmir Himalaya: progress, problems and prospects. Sains Malays 42(10):1377–1386.

Darwin C (1859) On the origin of species. published on, 24, 1.

David P, Thebault E, Anneville O, Duyck PF, Chapuis E, Loeuille N (2017) Impacts of invasive species on food webs: a review of empirical data. Adv Ecol Res 56:1–60.

De Terra H (1934) Physiographic results of a recent survey in Little Tibet. Geogr Rev 24(1):12–41.

Diagne C, Leroy B, Vaissière AC, Gozlan RE, Roiz, Jarić I et al (2021) High and rising economic costs of biological invasions worldwide. Nature 592(7855):571–576.

Downey PO, Richardson DM (2016) Alien plant invasions and native plant extinctions: a six-threshold framework. AoB plants, 8.

Duarte M, Verdú M, Cavieres LA, Bustamante RO (2021) Plant–plant facilitation increases with reduced phylogenetic relatedness along an elevation gradient. Oikos 130(2):248–259.

Eiserhardt WL, Svenning JC, Borchsenius F, Kristiansen T, Balslev H (2013) Separating environmental and geographical determinants of phylogenetic community structure in Amazonian palms (Arecaceae). Bot J Linn 171(1):244–259.

Elton CS (2020) The ecology of invasions by animals and plants. Springer Nature.

Enders M, Havemann F, Ruland F, Bernard Verdier M, Catford JA, Gómez-Aparicio L, et al (2020) A conceptual map of invasion biology: Integrating hypotheses into a consensus network. Glob Ecol Biogeogr 29(6):978–991.

Erneberg M (1999) Effects of herbivory and competition on an introduced plant in decline. Oecologia 118(2):203–209.

Essl F, Lenzner B, Bacher S, Bailey S, Capinha C, Daehler C (2020) Drivers of future alien species impacts: An expert[based assessment. Glob Chang Biol 26(9):4880–4893.

Gantchoff M, Wang G, Beyer D, Belant J (2018) Scale[dependent home range optimality for a solitary omnivore. Ecol Evol 8(23):12271–12282.

Gao GF, Peng D, Zhang Y, Li Y, Fan K, Tripathi BM et al (2021) Dramatic change of bacterial assembly process and co-occurrence pattern in Spartina alterniflora salt marsh along an inundation frequency gradient. Sci Total Environ 755:142546.

Godoy O, Kraft NJB, Levine J (2014) Phylogenetic relatedness and the determinants of competitive outcomes. Ecology Letters 17: 836–844.

González-Caro S, Umaña MN, Álvarez E, Stevenson PR, Swenson NG (2014) Phylogenetic alpha and beta diversity in tropical tree assemblages along regional-scale environmental gradients in northwest South America. J Plant Ecol 7(2):145–153.

Graham CH, Parra JL, Tinoco BA, Stiles FG, McGuire JA (2012) Untangling the influence of ecological and evolutionary factors on trait variation across hummingbird assemblages. Ecology 93(sp8):S99–S111.

Gupta SR, Singh JS (1982) Influence of floristic composition on the net primary production and dry matter turnover in a tropical grassland. Aust Ecol 7(4):363–374.

Hejda M, Pyšek P, Jarošík V (2009) Impact of invasive plants on the species richness, diversity and composition of invaded communities. J Ecol 97(3):393–403.

Hillebrand H, Bennett DM, Cadotte MW (2008) Consequences of dominance: a review of evenness effects on local and regional ecosystem processes. Ecol 89(6): 1510–1520.

IPBES (2019): Global assessment report on biodiversity and ecosystem services of the Intergovernmental Science-Policy Platform on Biodiversity and Ecosystem Services. ES Brondizio, J Settele, S Díaz, and HT Ngo (editors). IPBES secretariat, Bonn, Germany. 1148 pages. https://doi.org/10.5281/zenodo.3831673

Jackson ML 1973 “Chemical composition of soils”. In Chemistry of soils, Edited by: Bear, F E. 71–141. New Delhi: Oxford and IBH.

Jin Y, Qian H (2019) V. PhyloMaker: an R package that can generate very large phylogenies for vascular plants. Ecography 42(8):1353–1359.

Kay QON (1971) Biological flora of the British Isles: *Anthemis cotula* L. J Ecol 59(2): 623–636.

Keane RM, Crawley MJ (2002) Exotic plant invasions and the enemy release hypothesis. Trends Ecol Evol 17(4):164–170.

Kearney M, Porter W (2009) Mechanistic niche modelling: combining physiological and spatial data to predict species’ ranges. Ecol Lett 12(4):334–350.

Khuroo AA, Rashid I, Reshi Z, Dar GH, Wafai BA (2007) The alien flora of Kashmir Himalaya. Biol Inv 9(3):269–292.

Kraft NJ, Cornwell WK, Webb CO, Ackerly DD (2007) Trait evolution, community assembly, and the phylogenetic structure of ecological communities. The American Naturalist, 170(2): 271–283.

Kusumuto B, Villalobos F, Shiono T, Kubota Y (2019) Reconciling Darwin’s naturalization and pre[adaptation hypotheses: An inference from phylogenetic fields of exotic plants in Japan. J Biogeogr 46(11):2597–2608.

Leibold MA, Economo EP, Peres Neto P (2010) Metacommunity phylogenetics: separating the roles of environmental filters and historical biogeography. Ecology letters, 13(10): 1290–1299.

Lembrechts JJ, Lenoir J, Nuñez MA, Pauchard A, Geron C, Bussé G et al (2018) Microclimate variability in alpine ecosystems as stepping stones for non[native plant establishment above their current elevational limit. Ecography 41(6):900–909.

Li x.H, Zhu x.X, Niu y, Sun H (2014) Phylogenetic clustering and overdispersion for alpine plants along elevational gradient in the Hengduan Mountains Region, southwest China. J Syst Evol 52(3):280–288.

Linders TEW, Schaffner U, Eschen R, Abebe A, Choge SK, Nigatu L (2019) Direct and indirect effects of invasive species: Biodiversity loss is a major mechanism by which an invasive tree affects ecosystem functioning. J Ecol 107(6):2660–2672.

Loiola PP, de Bello F, Chytrý M, Götzenberger L, Carmona C P, Pyšek P, Lososová Z (2018) Invaders among locals: Alien species decrease phylogenetic and functional diversity while increasing dissimilarity among native community members. Journal of Ecology 106(6): 2230–2241.

Luo Y, Zhou M, Jin S, Wang Q, Yan D (2023) Changes in phylogenetic structure and species composition of woody plant communities across an elevational gradient in the southern Taihang Mountains, China. Global Ecology and Conservation, 42, e02412.

Mack MC (2003) Phylogenetic constraint, absent life forms, and preadapted alien plants: a prescription for biological invasions. International Journal of Plant Sciences, 164: S185– S196

Machac A, Janda M, Dunn RR, Sanders NJ (2011) Elevational gradients in phylogenetic structure of ant communities reveal the interplay of biotic and abiotic constraints on diversity. Ecography 34(3):364–371.

Manish K, Pandit MK (2018) Phylogenetic diversity, structure and diversification patterns of endemic plants along the elevational gradient in the Eastern Himalaya. Plant Ecol Divers 11(4):501–513.

McFadden IR, Sandel B, Tsirogiannis C, Morueta Holme N, Svenning JC, Enquist BJ, Kraft NJ (2019) Temperature shapes opposing latitudinal gradients of plant taxonomic and phylogenetic β diversity. Ecol Lett 22(7):1126–1135.

Miller ET, Farine DR, Trisos CH (2017) Phylogenetic community structure metrics and null models: a review with new methods and software. Ecography 40(4):461–77.

Ng J, Weaver WN, Laport RG (2019) Testing Darwin’s Naturalization Conundrum using phylogenetic relationships: Generalizable patterns across disparate communities? Divers Distrib 25(3):361–373.

Novotny V, Basset Y, Miller SE,Weiblen GD, Bremer B, Cizek L, Drodz P (2002) Low host specicity of herbivorous insects in a tropical forest. Nature 416: 841–844.

Olsen SR, Watanabe FS, Cosper HR, Larson WE, Nelson LB (1954) Residual phosphorus availability in long-time rotations on calcareous soils. Soil Sci 78(2):141–152.

Omer A, Fristoe T, Yang Q, Razanajatovo M, Weigelt P, Kreft H, … van Kleunen M (2022) The role of phylogenetic relatedness on alien plant success depends on the stage of invasion. Nature Plants 8(8): 906–914.

Paini DR, Sheppard AW, Cook DC, De Barro PJ, Worner SP, Thomas MB (2016) Global threat to agriculture from invasive species. Proc Natl Acad Sci 113(27):7575–7579.

Park DS, Feng X, Maitner BS, Ernst KC, Enquist BJ (2020) Darwin’s naturalization conundrum can be explained by spatial scale. Proc Natl Acad Sci 117(20):10904–10910.

Peay KG, Belisle M, Fukami T (2012) Phylogenetic relatedness predicts priority effects in nectar yeast communities. Proc Royal Soc B: Biol Sci 279(1729):749–758.

Pinto-Ledezma, JN, Larkin DJ, Cavender-Bares, J (2018) Patterns of beta diversity of vascular plants and their Correspondence with biome boundaries across North America. Front Ecol Evol 6:194.

Pinto-Ledezma, JN, Villalobos F, Reich PB, Catford JA, Larkin DJ, Cavender-Bares, J (2020) Testing Darwin’s naturalization conundrum based on taxonomic, phylogenetic, and functional dimensions of vascular plants. Ecol Monogr 90(4):e01420.

POWO (2021) Plants of the World Online. Facilitated by the Royal Botanic Gardens, Kew. Published on the Internet.

Pyšek P, Hulme PE, Simberloff D, Bacher S, Blackburn TM, Carlton J et al (2020) Scientists’ warning on invasive alien species. Biol Rev 95(6):1511–1534.

Qian H (2017) Climatic correlates of phylogenetic relatedness of woody angiosperms in forest communities along a tropical elevational gradient in South America. J Plant Ecol 11:394–400.

Qian H, Sandel B (2017) Phylogenetic relatedness of native and exotic plants along climate gradients in California, USA. Divers Distrib 23(11):1323–1333.

Qian H, Zhang Y, Zhang J, Wang X (2013) Latitudinal gradients in phylogenetic relatedness of angiosperm trees in North America. Glob Ecol Biogeogr 22(11):1183–1191.

R Core Team (2022). R: A language and environment for statistical computing. R Foundation for Statistical Computing, Vienna, Austria. URL https://www.R-project.org/.

Rejmánek M (1996) A theory of seed plant invasiveness: The first sketch. Biol Conserv 78: 171–181 doi:https://doi.org/10.1016/0006-3207(96)00026-2 (1996).

Rashid I, Reshi ZA (2012) Allelopathic interaction of an alien invasive specie Anthemis cotula on its neighbours Conyza canadensis and Galinsoga parviflora. Allelopathy J 29(1).

Rashid I, Reshi Z, Allaie RR, Wafai BA (2007) Germination ecology of invasive alien Anthemis cotula helps it synchronise its successful recruitment with favourable habitat conditions. Ann Appl Biol 150(3):361–369.

Reshi ZA, Shah MA, Rashid I, Rasool N (2012) Anthemis cotula L.: a highly invasive species in the Kashmir Himalaya, India. Invasive Alien Plants: an ecological appraisal for the Indian Subcontinent. CAB International, Oxfordshire, UK, 108-125.

Ricciardi A, Mottiar M (2006) Does Darwin’s naturalization hypothesis explain fish invasions? Biol Inv 8(6):1403–1407.

Richardson DM, Pyšek P (2012) Naturalization of introduced plants: ecological drivers of biogeographical patterns. New Phytol 196(2):383–396.

Rodgers WA, Panwar SH (1988) Biogeographical classification of India. New Forest, Dehra Dun, India.

Sander J, Wardell-Johnson G (2011) Fine[scale patterns of species and phylogenetic turnover in a global biodiversity hotspot: Implications for climate change vulnerability. J Veg Sci 22(5):766–780.

Sears MW, Raskin E, Angilletta Jr MJ (2011) The world is not flat: defining relevant thermal landscapes in the context of climate change. Integ Comp Biol 51(5):666–675.

Seebens H, Bacher S, Blackburn TM, Capinha C, Dawson W, Dullinger S et al (2021) Projecting the continental accumulation of alien species through to 2050. Glob Chang Biol 27(5):970–982.

Seebens H, Blackburn TM, Dyer EE, Genovesi P, Hulme PE, Jeschke J et al (2017) No saturation in the accumulation of alien species worldwide. Nat Commun 8(1):1–9.

Shah MA, Reshi Z (2007) Invasion by alien *Anthemis cotula* L. in a biodiversity hotspot: Release from native foes or relief from alien friends? Curr Sci 92(1):21–22.

Shah MA, Reshi ZA, Rashid I (2012) Synergistic effect of herbivory and mycorrhizal interactions on plant invasiveness. African Journal of Microbiology Research 6(19):4107–12.

Sheppard CS, Carboni M, Essl F, Seebens H, DivGrass Consortium, Thuiller W (2018) It takes one to know one: Similarity to resident alien species increases establishment success of new invaders. Diversity and Distributions, 24(5): 680–691.

Slingsby JA, Verboom GA (2006) Phylogenetic relatedness limits co-occurrence at fine spatial scales: evidence from the schoenoid sedges (Cyperaceae: Schoeneae) of the Cape Floristic Region, South Africa. Am Nat 168(1):14–27.

Smith SA, Brown JW (2018) Constructing a broadly inclusive seed plant phylogeny. Am J Bot 105(3):302–314.

Sol D, Garcia Porta J, González Lagos C, Pigot AL, Trisos C, Tobias JA (2022) A test of Darwin’s naturalization conundrum in birds reveals enhanced invasion success in the presence of close relatives. Ecology Letters, 25(3): 661–672.

Sommer B, Sampayo EM, Beger M, Harrison PL, Babcock RC, Pandolfi JM (2017) Local and regional controls of phylogenetic structure at the high-latitude range limits of corals. Proc Royal Soc B: Biol Sci 284(1861):20170915.

Thuiller W, Gallien L, Boulangeat I, De Bello F, Münkemüller T, Roquet C, Lavergne S (2010) Resolving Darwin’s naturalization conundrum: a quest for evidence. Divers Distrib 16(3):461–475.

Turbelin AJ, Malamud BD, Francis RA (2017) Mapping the global state of invasive alien species: patterns of invasion and policy responses. Global Ecol Biogeogr 26(1): 78–92.

Vetaas OR (1997) Relationships between floristic gradients in a primary succession. J Veg Sci 8(5):665–676.

Walkley A, Black IA (1934) An examination of the Degtjareff method for determining soil organic matter, and a proposed modification of the chromic acid titration method. Soil Sci 37(1):29–38.

Walsh JR, Carpenter SR, Vander Zanden MJ (2016) Invasive species triggers a massive loss of ecosystem services through a trophic cascade. Proc Natl Acad Sci 113(15):4081–4085.

Wang J, Li SP, Ge Y, Wang XY, Gao S, Chen T, Yu FH (2023) Darwin’s naturalization conundrum reconciled by changes of species interactions. Ecology, 104(1), e3850.

Webb CO, Ackerly DD, McPeek MA, Donoghue MJ (2002) Phylogenies and community ecology. Ann Rev Ecol Syst 475–505.

Webb CO, Losos JB, Agrawal AA (2006) Integrating phylogenies into community ecology. Ecol 87:S1–S2.

