## Supplementary figures and images for "Phylogenetic relatedness of plant species co-occurring with an invasive alien plant species (*Anthemis cotula* L.) varies with elevation"

### Fig. S4

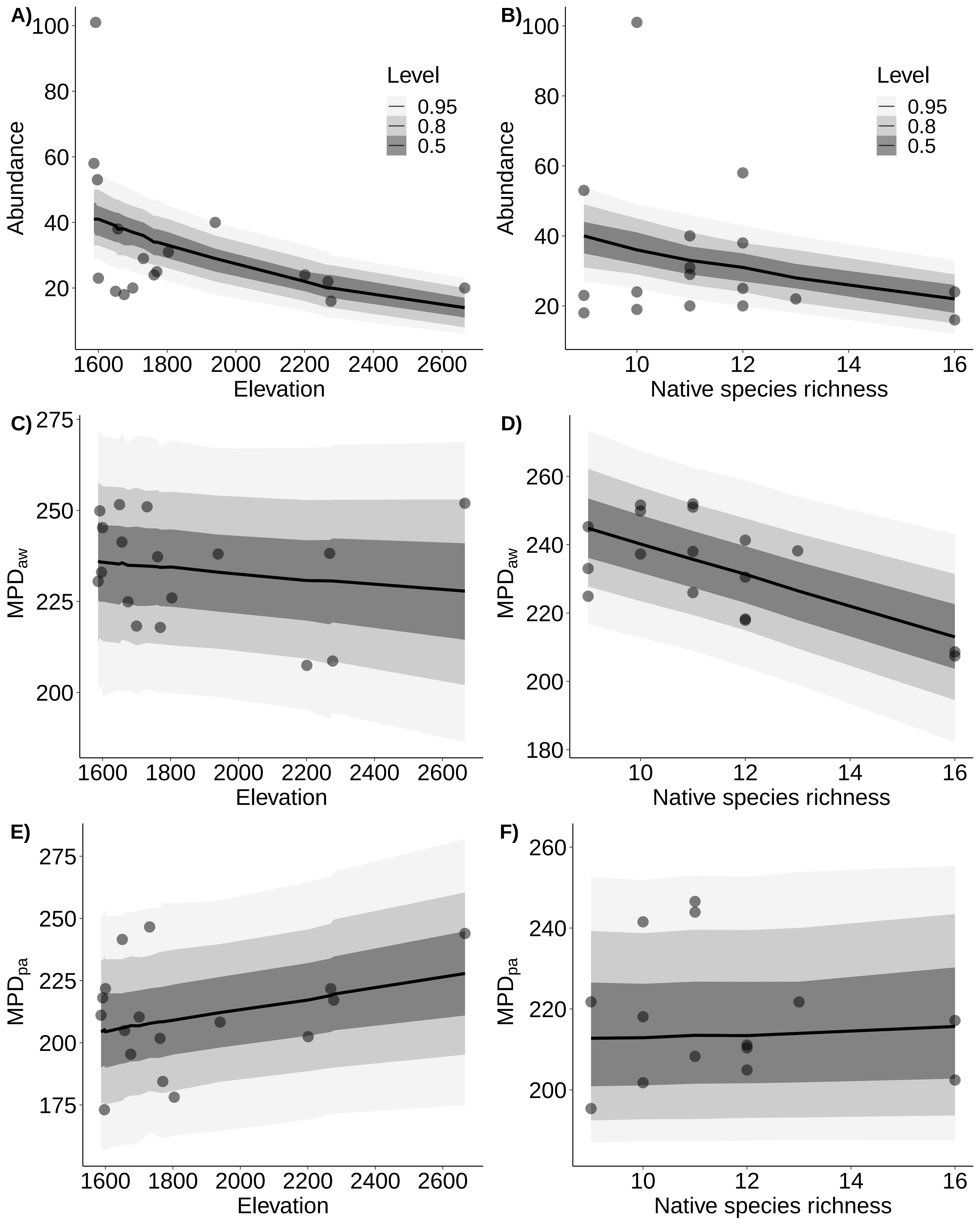
