## Supplementary material for "Phylogenetic relatedness of plant species co-occurring with an invasive alien plant species (*Anthemis cotula* L.) varies with elevation": Figures_S

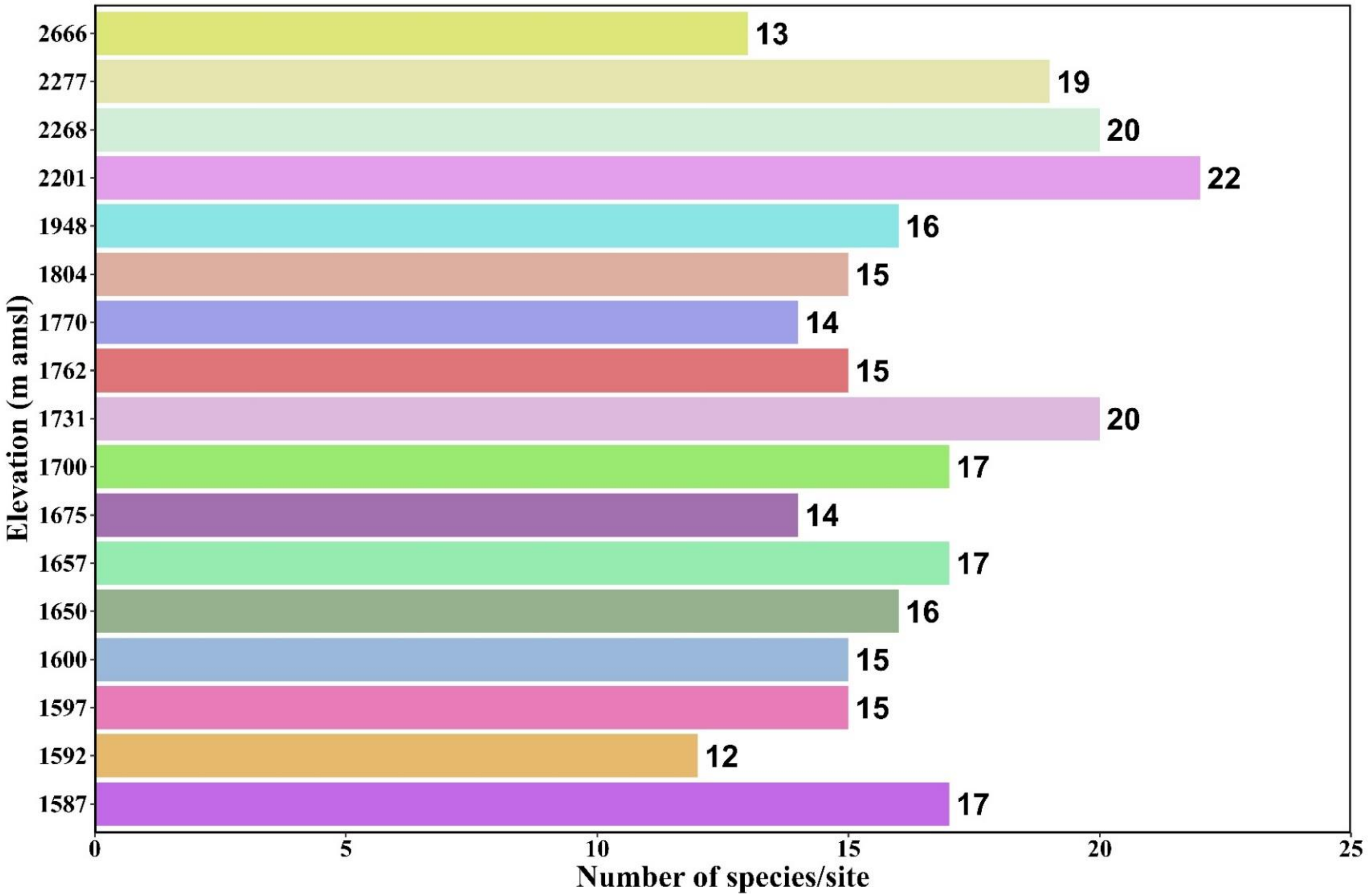


**Fig. S1**. Number of plant species across various sites at different elevations

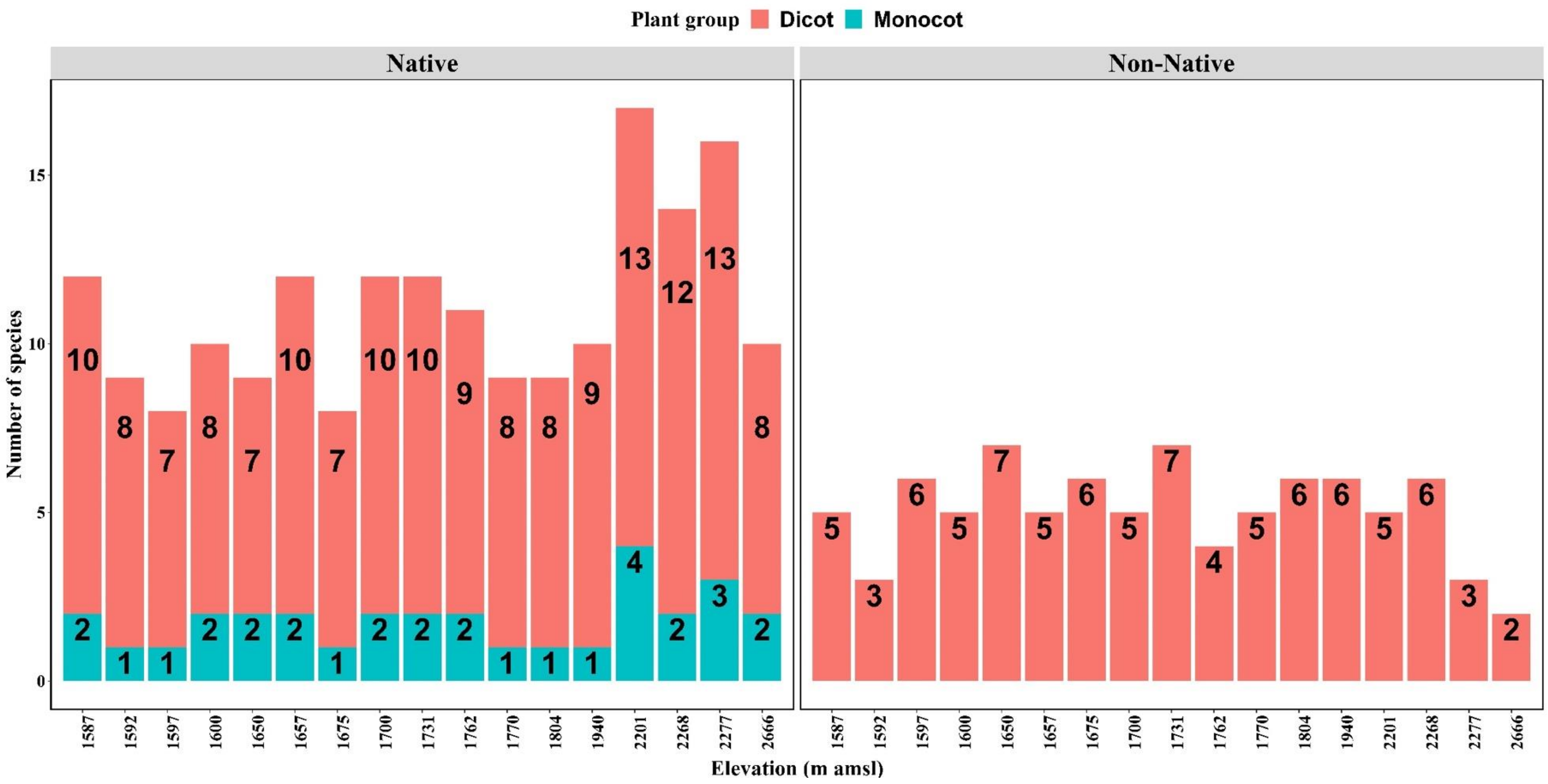


**Fig. S2.** Number of native and non-native species belonging to dicots and monocots associated with *Anthemis cotula* at different elevations.

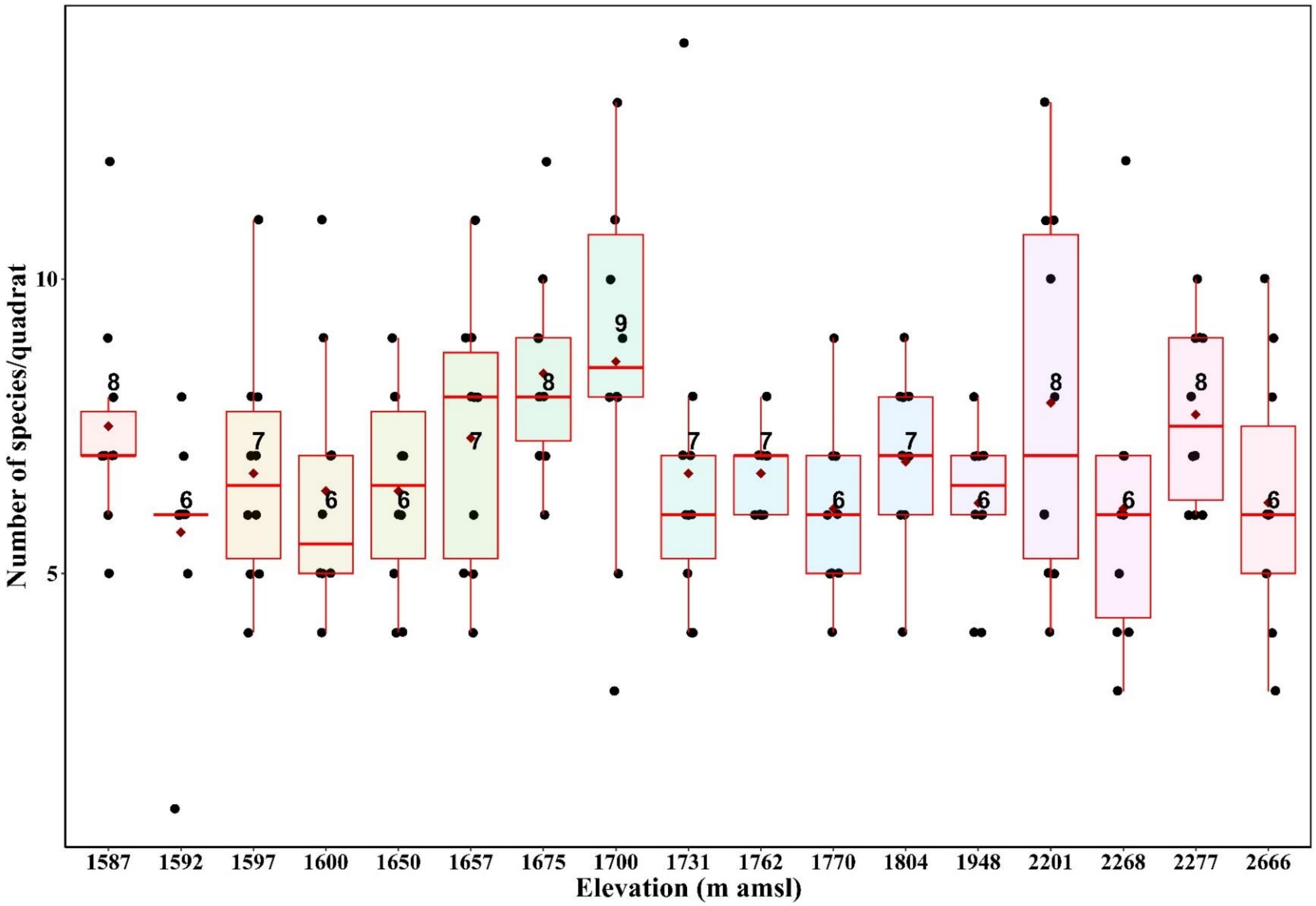


**Fig. S3.** Number of species per quadrat across sites at different elevations.

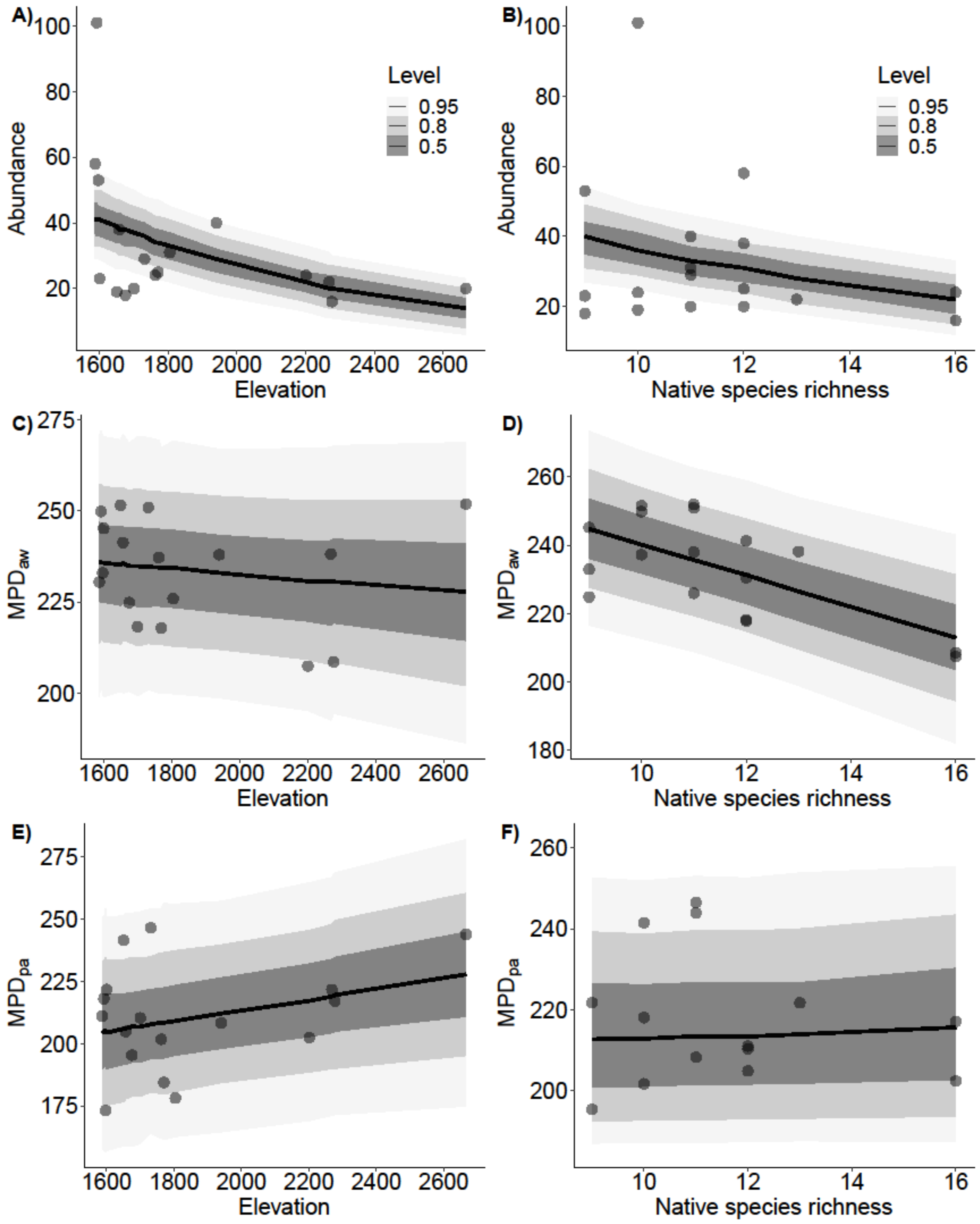


**Fig. S4.** Marginal effects plots of changes in abundance and phylogenetic structure of *A. cotula* across elevation (lefthand panels) and local species richness (righthand panels) for the natives-only dataset. Continuous black lines represent fitted slopes (with 50, 80, and 95% confidence intervals in gray).
