## Supplementary material for "Phylogenetic relatedness of plant species co-occurring with an invasive alien plant species (*Anthemis cotula* L.) varies with elevation": Table_S1

Table S1. Geographical coordinates, elevation and soil characteristics of study sites

| **Sites** | **Elevation (mamsl)** | **Latitude** | **Longitude** | **pH** | **SOC (%)** | **N (ppm)** | **P (ppm)** | **K (ppm)** | **S (ppm)** | **Ca (ppm)** |
| --- | --- | --- | --- | --- | --- | --- | --- | --- | --- | --- |
| Site 1 | 1587 | 34˚13’56” N | 74˚43’27”E | 8 | 1.48 | 83.7 | 2.23 | 66.96 | 2.92 | 1469 |
| Site 2 | 1592 | 34˚07̇̍’47”N | 74˚50’16”E | 8.4 | 2.1 | 89.28 | 4.01 | 61.38 | 4.3 | 1924 |
| Site 3 | 1597 | 34˚25’23”N | 74˚38’02”E | 7.2 | 2.73 | 92.07 | 3.57 | 59.15 | 2.7 | 1316 |
| Site 4 | 1600 | 33˚37’51”N | 75˚14’2”E | 7.54 | 3.31 | 103.23 | 2.67 | 54.68 | 2.5 | 1611 |
| Site 5 | 1650 | 34˚02’10”N | 74˚40’33”E | 7.8 | 2.31 | 80.35 | 3.125 | 62.72 | 2.93 | 1779 |
| Site 6 | 1657 | 33˚52’13”N | 74˚53’41”E | 7.7 | 2.54 | 92.52 | 3.125 | 64.06 | 2.5 | 1648 |
| Site 7 | 1675 | 34˚20’03”N | 74˚40’50”E | 7.9 | 2.75 | 76.11 | 3.125 | 65.1 | 3.02 | 1395 |
| Site 8 | 1700 | 34˚30’55”N | 74˚10’22”E | 7.6 | 2.92 | 111.6 | 3.57 | 64.73 | 3.32 | 1222 |
| Site 9 | 1731 | 34˚10’12”N | 74˚28’17”E | 7.4 | 2.73 | 92.07 | 2.67 | 65.84 | 2.8 | 1663 |
| Site 10 | 1762 | 33˚37’31”N | 75˚14’05”E | 7.1 | 2.01 | 125.55 | 4.01 | 64.73 | 3.75 | 1478 |
| Site 11 | 1770 | 34˚00’45”N | 74˚35’55”E | 7 | 2.02 | 129.46 | 2.67 | 63.61 | 2.95 | 1745 |
| Site 12 | 1804 | 34˚15’35”N | 74˚54’24”E | 7.12 | 3.65 | 92.07 | 3.57 | 63.61 | 2.73 | 1386 |
| Site 13 | 1948 | 33˚34’29”N | 75˚19’17”E | 7.09 | 3.54 | 102.67 | 3.57 | 53.68 | 2.56 | 1389 |
| Site 14 | 2201 | 34˚16’01”N | 75˚08’54”E | 5.9 | 3.14 | 89.28 | 3.57 | 64.95 | 3.12 | 1424 |
| Site 15 | 2268 | 34˚16’28”N | 75˚10’51”E | 6.32 | 4.45 | 84.93 | 3.125 | 58.03 | 2.14 | 1428 |
| Site 16 | 2277 | 33˚40’59”N | 74˚47’34”E | 6.9 | 3.56 | 112.61 | 3.125 | 61.6 | 2.6 | 1318 |
| Site 17 | 2666 | 33˚39’41”N | 74˚47’04”E | 6.36 | 5.32 | 83.92 | 3.125 | 64.06 | 2.63 | 1445 |
